# Lung influenza virus specific memory CD4 T cell location and optimal cytokine production are dependent on interactions with lung antigen-presenting cells

**DOI:** 10.1101/2023.09.19.558387

**Authors:** Kerrie E Hargrave, Julie C Worrell, Chiara Pirillo, Euan Brennan, Andreu Masdefiol Garriga, Joshua I Gray, Thomas Purnell, Edward W Roberts, Megan KL MacLeod

## Abstract

Influenza A virus (IAV) infection leads to the formation of mucosal memory CD4 T cells that can protect the host. An in-depth understanding of the signals that shape memory cell development is required for more effective vaccine design. We have examined the formation of memory CD4 T cells in the lung following IAV infection of mice, characterising changes to the lung landscape and immune cell composition. IAV-specific CD4 T cells were found throughout the lung at both primary and memory time points. These cells were found near lung airways and in close contact with a range of immune cells including macrophages, dendritic cells, and B cells. Interactions between lung IAV-specific CD4 T cells and MHCII+ cells during the primary immune response were important in shaping the subsequent memory pool. Treatment with an anti-MHCII blocking antibody increased the proportion of memory CD4 T cells found at lung airways but reduced interferon-g expression by IAV-specific immunodominant memory CD4 T cells. The immunodominant CD4 T cells expressed higher levels of PD1 than other IAV-specific CD4 T cells and PD1+ memory CD4 T cells were located further away from MHCII+ cells than their PD1-negative counterparts. This distinction in location was lost in mice treated with anti-MHCII antibody. These data suggest that sustained antigen presentation in the lung impacts on the formation of memory CD4 T cells by regulating their cytokine production and location.

## 1 Introduction

Influenza A (IAV) virus is a common pathogen that infects the respiratory tract. Despite the availability of vaccines, IAV continues to present a substantial global health challenge. Current vaccines generate an antibody response that targets the highly variable surface proteins of the virus. While antibodies can neutralise the virus, seasonal viral mutation and immune selection make it challenging for antibodies to recognise modified viruses^1, 2^. The leads to viral immune evasion, the outbreak of seasonal epidemics, and the risk of pandemics.

In contrast to antibodies, CD4 T cells often recognise viral regions conserved between different IAV strains^3^. Following IAV infection, activated CD4 T cells migrate to the lung where they play diverse roles in viral control including helping CD8 T cells and B cells, in addition to mediating direct effector functions via cytokines^4, 5^. Some of these recruited CD4 T cells will remain in the lung after infection as tissue resident memory (Trm) cells. Lung memory CD4 T cells can provide enhanced protection from re-infection compared to memory cells from secondary lymphoid tissues by acting directly and through interactions with other cells to limit viral replication^6–8^. Importantly, the presence of IAV specific CD4 T cells correlates with reduced disease symptoms in humans^9, 10^.

Typically, Trm cells are examined by flow cytometry based on their persistence within the tissue, expression of certain retention molecules, and lack of binding to antibodies injected into the blood shortly before analysis^11, 12^. While there is a growing understanding of the heterogeneity of Trm cells,^13, 14^ much less is known about the location of Trm within the lung and whether this alters following resolution of the infection. Here we have used reporter mice to identify lung IAV-specific CD4 T cells at multiple time points following infection. We identified IAV-specific CD4 T cells dispersed throughout the lung, at airways, and within clusters with other immune cells, including various types of MHCII+ antigen presenting cells (APCs). While the composition of the APCs altered during the infection time course, CD4 T cells were found in close proximity to MHCII+ cells at both primary and memory time points.

The close proximity between IAV-specific CD4 T cells and these APCs indicates a role for antigen presentation in the generation of memory T cells. Inhibiting the interactions between CD4 T cells and APCs with a blocking anti-MHCII antibody can disrupt early T cell responses and/or the formation of immune memory depending on the timing of delivery^15–17^. Here we have delivered anti-MHCII intranasally following the recruitment of CD4 T cells to the lung to investigate the role for interactions between lung CD4 T cells and APCs in the generation of immune memory.

We found that the numbers of IAV specific memory CD4 T cells were not altered by anti-MHCII treatment. However, the location of IAV-specific memory CD4 T cells and their ability to produce the key anti-viral cytokine, interferon (IFN)g, were altered by the blockade. We found that anti-MHCII treatment increased the percentage of memory cells found at airways but reduced IFNg production by CD4 T cells that recognise an immunodominant IAV nucleoprotein (NP) epitope. Together, these data indicate that interactions between lung APCs and CD4 T cells influence the quality and location of memory CD4 T cells generated by infection, findings that have important implications for the design of mucosal vaccines.

## 2. Materials and Methods

### 2.1 Animals

10-week-old female C57BL/6 mice were purchased from Envigo (UK) and male and female TRACE mice were bred at the University of Glasgow. TRACE and C57BL/6 mice were maintained at the University of Glasgow under specific pathogen free conditions in accordance with UK home office regulations (Project Licenses P2F28B003 and PP1902420) and approved by the local ethics committee. TRACE mice have been described previously^18, 19^.

### 2.2 Influenza A virus (IAV) Infections

TRACE or C57BL/6 mice were briefly anesthetised using inhaled isoflurane and infected with 100-200 plaque forming units of IAV WSN strain in 20ul of PBS intranasally (i.n.) depending on their age, weight, and sex. Influenza A virus (IAV) was prepared and titered in MDCK cells. Infected mice were weighed daily from day 4 post-infection. Any animals that lost more than 20% of their starting weight were humanely euthanised. TRACE mice were given Dox+ chow (Envigo) for a total of 12 days starting two days prior to infection.

### 2.3 Intranasal antibody instillation

Mice were anesthetised using inhaled isoflurane at days 6 and 12 post-infection and 100mg of anti-MHCII (Y3P) or rat IgG2a (2A3), both from BioXcell, instilled in 30ml of PBS. For Y3P labelled studies, Y3P and IgG were labelled with Alexa-Fluor 546 (ThermoFisher) according to the manufacturer’s instructions. Mice received the labelled antibody on day 6 post-infection and were euthanised after 2 hours.

### 2.4 Digital Slide scanning

Excised lung tissues for histological analysis were fixed in 10% neutral buffered formalin for 24 hours and paraffin embedded in a tissue processor (Shandon Pathcentre® Tissue Processor). Formalin-fixed and paraffin-embedded (FFPE) tissues were sectioned (6μm) and processed for subsequent staining with Hematoxylin and Eosin (H&E) using standard protocols. Stained slides were scanned on a Leica Aperio VERSA 8 bright-field slide scanner (Leica Biosystems) at 10X magnification and were visualised/analysed using Aperio ImageScope version 12.1.0.5029 (Aperio Technologies Inc., Vista).

### 2.5 Blinded analysis of H&E staining

Manual identification of selected regions of interest (ROIs) was performed by a blinded observer. ImageScope software (Aperio Technologies Inc) was used to provide independent outputs for each tissue section. This included quantifying the number of areas of high cell density (haematoxylin stained, clusters), measurement of the cluster area (expressed as µm^2^) measurement of distance/proximity between clusters and the nearest airway (expressed as µm) and assessment of damaged area (expressed as % of total lung area). The Aperio nuclear V9 algorithm (Aperio ScanScope XT System; Aperio Technologies) was used to count the number of cells contained within a cluster. Damaged areas were identified as regions of interest that were not airways, blood vessels, immune cell clusters or normal /healthy lung tissue as found in the naïve animals.

### 2.6 Immunofluorescent imaging

Mice were euthanised by rising concentrations of CO_2_ and the vena cava cut. Lungs were perfused with PBS-5mM Ethylenediaminetetraacetic acid (EDTA) to remove red blood cells and 1% paraformaldehyde (PFA) to fix the lungs and preserve the integrity of the EYFP protein. Lung inflation was achieved using 1-3ml 1% warm UtraPure low melt agarose administrated via the trachea. Lungs with solidified agarose were incubated in 1% PFA overnight at 4°C followed by incubation in 30% sucrose for a further 2-5 days at 4°C. Lung lobes were frozen in OCT (Tissue-Tek 4583) and stored at -80°C. Lungs were sectioned into 10um slices on a Shandon Cryotome FE (Thermo Scientific 12087159) and mounted onto Super Frost slides (Thermo Scientific). Slides were fixed in 100% cold acetone and stored at -20°C.

Slides were rehydrated in PBS containing 0.5% bovine serum albumin (BSA) for 5 minutes and incubated with Fc Block (24G2) for 30-60 minutes at room temperature. The sections were stained with antibodies at 4°C overnight: anti-CD4 AlexaFluor647 (BD, RM4-5), anti-MHCII eFluor 450 (M5114, ThermoFisher), anti-B220-PE (ThermoFisher, RA3-6B2), anti-CD64-PE (BioLegend, X54-5/7.1), anti-CD11c-Alexa-594 (BioLegend, N418), anti-PD1-PE (BioLegend, 29F.1A12), anti-GFP-Alexa 488 (rabbit polyclonal, Invitrogen). Slides were washed in PBS-BSA and mounted with VectorSheild (Vector). Images were collected on a LSM880 (Ziess) confocal microscope at 20X magnification.

For PCLS, lungs inflated with agarose were sectioned at 300mm using a vibrotome (5100mz Campden Instruments) and stored at 4°C in PBS containing 1% BSA and 0.05% sodium azide. Sections were incubated with protein block (Abcam) before the addition of primary antibodies: anti-CD4 AlexaFluor647 (BD, RM4-5), anti-MHCII eFluor 450 (M5114, ThermoFisher), anti-CD68 AF594 (FA-11, BioLegend) and anti-GFP-Alexa 488 (rabbit polyclonal, Invitrogen) diluted in antibody diluent reagent solution (Life technologies). The slices were incubated overnight at 4°C and then washed with wash buffer (1% BSA, 0.1% TritonX-100 and 0.05% sodium azide in PBS). Ce3D clearing solution (Biolegend) was added 30 minutes before mounting in a seal frame incubation chamber (ThermoFisher) and covered with a coverslip. Images were acquired using a Zeiss LSM 880 Airyscan confocal microscope and analysed using Imaris software.

### 2.7 Image analysis

Confocal slide images were prepared in Zen (Ziess) and analysed in Volocity (Version 6, Quorum Technologies). Objects were defined based on fluorescence, object size and pixels, in cases of high cell density, touching objects were separated. Cells were manually checked when used for measurement analysis. Distances were measured using the measuring tool and regions of interests around airways cropped to count cells at airways.

### 2.8 Tissue preparation for CD4 T cell analysis by flow cytometry

Prior to euthanasia by cervical dislocation, TRACE mice were injected intravenously (i.v.) with 1mg PE conjugated anti-CD45 (clone; 30F11) (ThermoFisher) for 3 minutes. Alternatively, in the C57BL/6 mouse experiments in Figure 5, mice were euthanised using a rising concentration of CO_2_ and perfused with PBS with 5mM EDTA through the heart. In both cases, single cell suspensions of lungs were prepared by digestion of snipped lung tissue with 1mg/ml collagenase and 30mg/ml DNAse (Sigma) for 40 minutes at 37°C in a shaking incubator and tissues disrupted by passing through a 100mm filter. Spleen and mediastinal lymph nodes were processed by mechanical disruption. Red blood cells were lysed from spleen and lungs with lysis buffer (ThermoFisher). Cells were counted using a haemocytometer with dead cells excluded using Trypan Blue.

### 2.9 Tissue preparation for APC analysis by flow cytometry

Mice were euthanised by cervical dislocation and lungs digested with a final concentration of 1.6mg/ml Dispase, 0.2mg/ml Collagenase P (Roche) and 0.1mg/ml DNase (Sigma) for 40 minutes at 37°C in a shaking incubator and tissues disrupted by passing through a 100mm filter. Red blood cells were lysed. Cells were counted using a haemocytometer with dead cells excluded using Trypan Blue.

### 2.10 Production of bone marrow derived dendritic cells

Bone marrow derived dendritic cells were generated from femurs and tibias of 8 -10week old C57BL/6 mice with RPMI 1640 (supplemented with 10% heat inactivated foetal calf serum, 100mg/ ml penicillin-streptomycin and 2mM L-glutamine). Cells were cultured at 37°C 5% CO_2_ in the presence of GM-CSF (conditioned from the supernatant of X63 cells^20^) for 7 days. Media was supplemented or replaced on day 2 and 5. After this time, adherent cells were harvested, seeded, and then incubated overnight with IAV antigen (MOI of 0.3) produced as described previously^21^.

### 2.11 Ex vivo reactivation for cytokine production

Lung single cell suspensions were co-cultured with IAV-Antigen+ bmDCs in complete RMPI at a ratio of approximately 10:1 T cells to DCs. Alternatively, cells were cultured with 10mg/ml NP peptide (QVYSLIRPNENPAHK, JPT). *Ex vivo* cultures were incubated at 37°C, 5% CO_2_ for 6 hours in the presence of Golgi Plug (BD Bioscience).

### 2.12 Flow cytometry staining

#### IAV-specific CD4 T cells

When required, single cell suspensions were stained with PE or APC-labelled IA^b^/NP_311-325_ tetramers (NIH tetramer core) for 2 hours at 37°C, 5% CO_2_ in complete RPMI containing Fc block (24G2). After this time, surface antibodies were added and the cells incubated for a further 20 minutes at 4°C. Alternatively, cells were incubated for 10 minutes at 4°C with Fcblock and surface antibodies added for a further 20 minutes. Antibodies used were: anti-CD4 APC-Alexa647 (RM4-5, ThermoFisher), anti-CD8 BUV805 (BD 53-6.7) or anti-CD8-eFluor 450 (ThermoFisher 53-6.7), anti-CD44 PerCP-Cy5.5 or BUV395 (ThermoFisher or BD: IM7), anti-PD1 BV605 (BioLegend 29F.1A12), anti-ICOS PerCP-Cy5.5 (ThermoFisher 7E.17G9), anti-B220 eFluor 450 (RA3-6B2), anti-MHCII eFluor 450 (M5114) and anti-F480 eFluor 450 (BM8), all ThermoFisher. Anti-B220, MHCII and F480, and CD8, were used as a ‘dump’ gate. Cells were stained with a fixable viability dye eFluor 506/Lyme (ThermoFisher).

For some experiments, cells were fixed with cytofix/cytoperm (BD Bioscience) for 20 minutes at 4°C and stained in perm wash buffer with anti-IFNg PE or BV785 (ThermoFisher or BioLegend, clone: XMG1.2) for one hour at room temperature.

#### Lung APCs

Cells were first labelled with Fc block for 10 minutes at 4°C and then the following antibodies added: CD45-PE (BD, 30-F.11), Siglec F-APC (BioLegend, S17007L), CD11b-PeCy7 (BioLegend, M1/70), Ly6G-BV785 (BioLegend, 1A8), Ly6C-PerCP-Cy5.5 (ThermoFisher, HK1.4), MHCII BUV395 (BD, 2G9), CD64-BV711 (BioLegend, X54-5/7.1), CD11c-eFluor780 (ThermoFisher, N418), B220-eFluor 450 (RA3-6B2, ThermoFisher), CD103-Qdot 605 (BioLegend, 2E7). Cells were then stained with a fixable viability dye eFluor 506/Lyme (ThermoFisher).

Stained cells washed with FACS buffer or Permwash following intracellular staining and acquired on a BD Fortessa and analysed using FlowJo (version 10).

### 2.13 Statist**i**cal analysis

Data were analysed using Prism version 10 (GraphPad). Differences between groups were analysed as indicated in figure legends depending on whether data were normally distributed. In all figures * represents a p value of <0.05; **: p>0.01, ***: p>0.001, ****: p>0.0001.

## 3. Results

### 3.1 IAV infection leads to sustained changes in the lung

To first characterise how influenza virus infection alters the landscape of the lung, we performed a blinded analysis of Haematoxylin and Eosin (H&E) stained lung sections taken at day 9 and 30 following infection with IAV (strain WSN), Figure 1A. H&E-stained sections were digitally scanned, and a blinded analysis was performed to identify IAV-induced alterations in the lung. Lungs from infected mice had clear areas of cell infiltrate including dense cell clusters, often found near airways and that are similar to structures described by others^22–28^.

**Figure 1.**
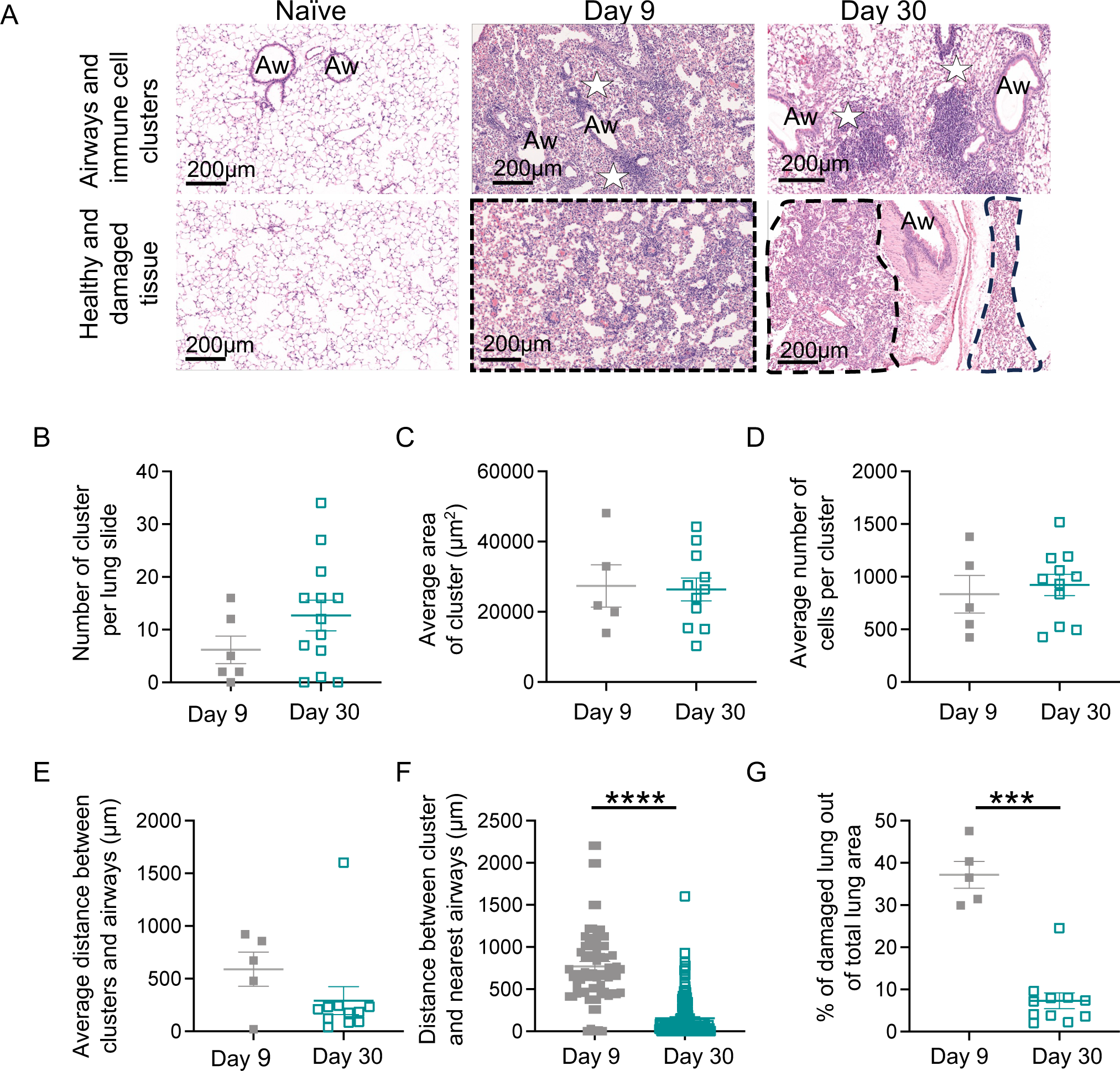
Persistent changes to the lung landscape following IAV infection. C57BL/6 mice were infected intranasally with IAV and lungs taken from naïve animals or at 9 or 30 days post-infection. A: H&E staining on lung sections from naïve and IAV infected mice; all images taken at 10x magnification, scale bar 200µm. Airways (Aw), immune cell clusters (white stars) and areas of damage (black dashed areas) are indicated. Data are representative of 6-13 mice per experimental group, each slide contained one section of each of the five lung lobes and data shown are added from across all lobes to provide one value per slide. Blinded analysis of lung sections was performed using ImageScope software to determine: B: the number of areas of high cell density clusters; C: average measurement of the cluster area (µm^2^); D: average number of cells contained within a cluster; E: average and F: individual distance/proximity between clusters and the nearest airway (µm); G: percentages of damaged lung areas; in C-F, 2 day 30 samples were removed as no clusters were present. In B-E, & G each symbol represents a mouse and the line is the mean of the group. In F, each symbol represents a cluster, and the line shows the mean of all clusters. In all figures, error bars are SEM. In F and G data are not normally distributed (tested by Shapiro-Wilk) and were analysed by Mann Whitney U test, ***: p<0.001, ****: p <0.0001.

We determined the number of clusters per lung, their size, cell number, and distance from the nearest airway. We identified an average of 6.2 clusters per lung section at day 9 and 12.7 at day 30 post-infection, Figure 1B. These data demonstrate that clusters are maintained despite viral clearance and formation of immune memory^29–31^. The cluster area and number of cells within the clusters also did not alter between the primary and memory time points, Figure 1C-D. However, the clusters tended to be closer to airways at memory time points. This difference was significant when the data were analysed as individual clusters, but not when we calculated an average per slide, Figure 1E-F. At both primary and memory time points, tissue sections contained areas of damage, often located near immune cell clusters. To calculate a percentage of the lung that contained such tissue, areas of damage that were not clearly organised cell clusters were expressed as a percentage of total lung area, Figure 1G. This measure reduced from the primary to the memory time point, consistent with repair to the lung following viral clearance^32^.

### 3.2 Immune cells are found throughout the lung, near airways and in cell clusters at primary and memory time points following IAV infection

To investigate the location of immune cells within the lung parenchyma, airways and clusters, we performed immunofluorescence analysis of lungs at days 10 and 40 post-infection. To first provide an overview of the lung immune response, we combined precision cut lung slices (PCLS) with tissue clearing and confocal microscopy. To enable us to identify IAV-specific CD4 T cells, we took advantage of a reporter mouse, TRACE, that we have developed and previously characterised^18, 19^. Activated T cells in these triple transgenic mice express rtTA under the control of part of the interleukin 2 promoter. When bound to doxycycline, rtTA activates the tet-ON promoter driving Cre expression. This leads to the irreversible labelling of activated T cells by expression of the fluorescent protein EYFP from the Rosa locus, Figure 2A. Infection with IAV induces a robust population of EYFP+ CD4 T cells when reporter mice are fed doxycycline in their diet during the initial infection^19^. Our prior studies have shown that EYFP+ cells are found in lung and secondary lymphoid organs following IAV infection^19^. This model greatly increases the number of responding T cells we can identify by both flow cytometry and immunofluorescence in comparison to other methodologies such as TCR transgenic cells or MHC tetramers. Moreover, it enables us to identify a broad range of responding CD4 T cells rather than identifying cells that respond to a single epitope.

**Figure 2.**
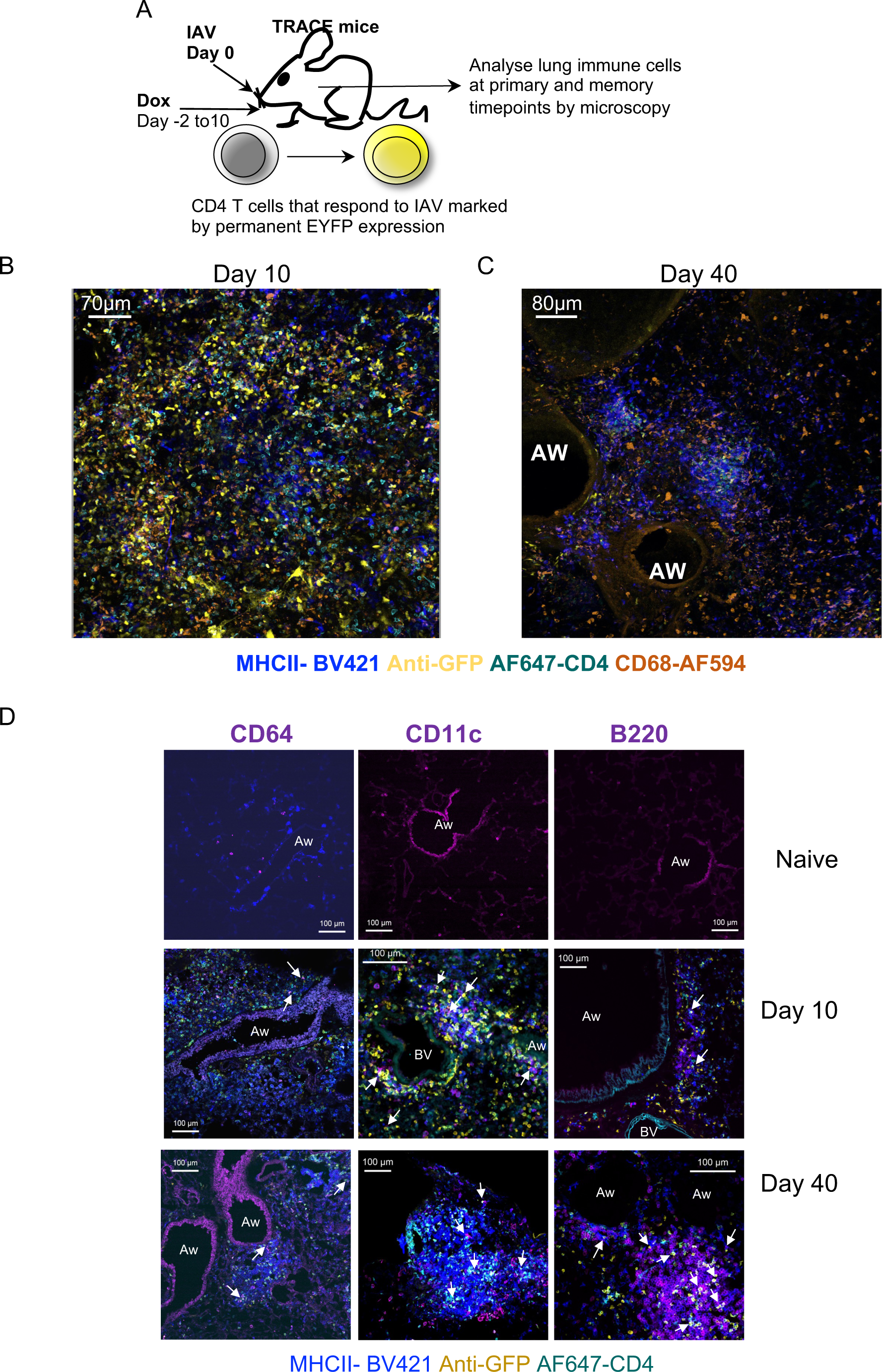
Immune cells are found throughout the lung, near airways and in cell clusters at primary and memory time points following IAV infection. A: TRACE mice enable identification of responding CD4 T cells via permanent expression of EYFP. Lungs from naïve or IAV infected TRACE mice were analysed 10 or 40 days post-IAV infection. In B-C, PCLS were cleared and stained with the indicated antibodies, scale bars are 70 (A) or 80 (B)mm. In D, frozen lungs were sectioned and stained with the indicated antibodies to identify IAV specific CD4 T cells, and MHCII+ CD64, CD11c or B220+ cells. Data are representative of one image set (B-C) or 3-5 mice per time point (D). White arrows indicate EYFP+CD4+ cells in close proximity to CD64, CD11c or B220+ cells, scale bars are 100mm.

The cleared lung tissue showed that multiple immune cell populations were distributed throughout the parenchyma at day 10 post-infection, Figure 2B. In contrast, at day 40 post-infection, immune cells were more obviously packed together in clusters, often around airways and similar to the structures we observed in the H&E analysis, Figure 2C.

We next used confocal microscopy of lung sections to more closely examine the cells within the clusters and the cells located near airways. A large number of cells were MHCII+ and we stained sections with antibodies to CD64, B220 and CD11c to provide insights into the identity of these cells, Figure 2D and Supplementary Figure 1. MHCII+ cells that also expressed either CD64, B220 and CD11c were identified in close proximity to IAV-specific CD4 T cells at both time points. The representative images demonstrate the diversity of the anti-viral response with some images illustrating tight networks of immune cells clustered together while others showing immune cells forming a line around airways.

As multiple cell types can express CD64, CD11c and B220, we used flow cytometry to identify and characterise different MHCII+ subsets in naïve animals and at primary and memory IAV infection time points. We first used the gating strategy in Supplementary Figure 2A to define multiple populations of myeloid or lymphoid cells and assessed the changes in the proportion of these cells across the time points. We then gated on MHCII+ cells that were either CD64+, B220+ or CD11c+, Supplementary Figure 2B, and overlaid the initial gates to identify which cell types were within these MHCII+ populations.

As expected^33, 34^, during the initial response there was an increase in monocyte and macrophage populations in the lung, Figure 3A. There were fewer lung conventional (c) DCs at day 6 post-infection compared to in naïve animals, indicating the migration of these cells to the draining lymph node^35, 36^. Most of these populations returned to levels found in naïve animals at the memory time point. However, we did find a sustained increase in the numbers of B cells and conventional (c)DC1 at day 30.

**Figure 3.**
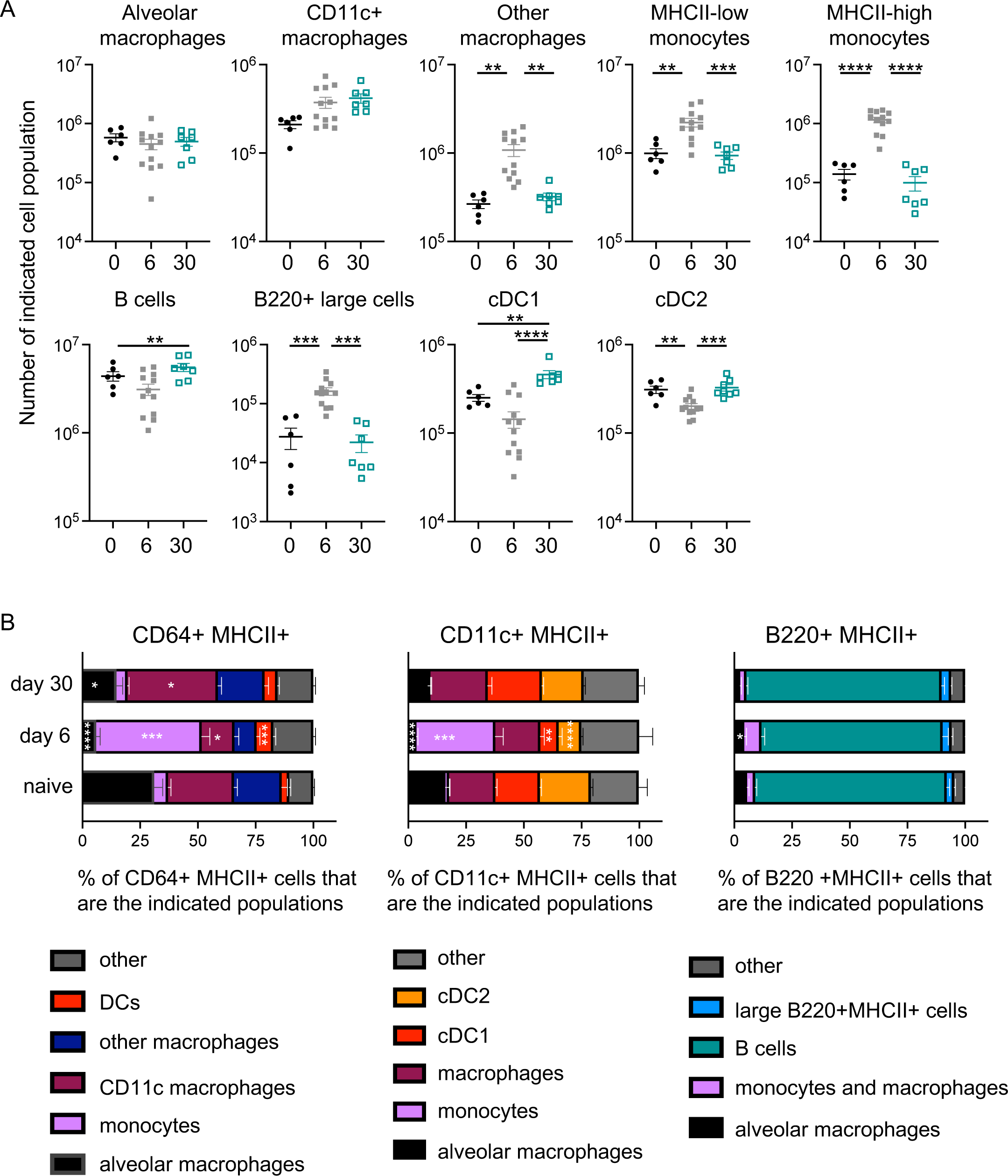
Lung MHCII+ populations alter across the IAV infection time course. C57BL/6 mice were infected with IAV and lungs taken at 6 or 30 days post-infection or from naïve animals. Following tissue digest, lungs were analysed by flow cytometry as indicated in Supplementary Figure 2. In A, each symbol represents a mouse and the line shows the mean of the group. Data are from 2 independent experiments per time point with a total of 6-12 per group. All data were normally distributed (tested by Shapiro-Wilk) and differences between time points assessed by ANOVA followed by a Tukey’s test. In B, cells were gated on MHCII+ cells that were CD64, CD11c or B220+ and the proportion of the indicated cell populations determined. Not all data were normally distributed and differences between infection time points and naïve samples tested by ANOVA followed by a Kruskal-Wallis test. In all plots, error bars are SEM and *: p<0.05; **:p<0.01; ***: p<0.001; ****p<0.0001.

In line with these changes at the cell population level, CD11c+MHCII+ cells and CD64+MHCII+ cells contained a greater proportion of monocytes and macrophages at day 6 post-infection Figure 3B. In these analyses, we have combined related populations, e.g. different macrophage populations, when one or both populations were present at low frequencies. By day 30, the proportional make-up of these populations was similar to that in naïve animals although the CD64+MHCII+ population contained slightly fewer alveolar macrophages and slightly more CD11c+ macrophages at day 30 compared to naïve animals. In contrast, the populations of B220+MHCII+ cells were consistent across the time course and were mainly B cells. Together, these data suggest that multiple different types of MHCII+ cells have the potential to interact with CD4 T cells in the lung during and following IAV infection.

### 3.3 IAV-specific CD4 T cells are found at the airway at primary and memory time points

CD4 T cells are known to persist in the lung following IAV infection^4, 5^ and we have characterised the number, phenotype and cytokine production by IAV specific CD4 T cells in previous studies^18, 19, 21, 37^. Similar to the MHCII+ cells, IAV-specific CD4 T cells were found in the lung parenchyma, at airways, and in immune clusters at primary and memory time points, Figures 2 and 4A. As expected, the numbers of EYFP+CD4+ cells found per slide declined from day 10 to day 40 post-infection, Figure 4B. Despite this, the percentages of these cells that were found at the airways were consistent between the two time points suggesting that memory CD4 T cells are just as likely to patrol the sites of infection during and beyond an active infection, Figure 4C. In contrast, the numbers of MHCII+ cells and the percentage of these cells found at airways did not change between the primary and memory time points, Figure 4D-E.

**Figure 4.**
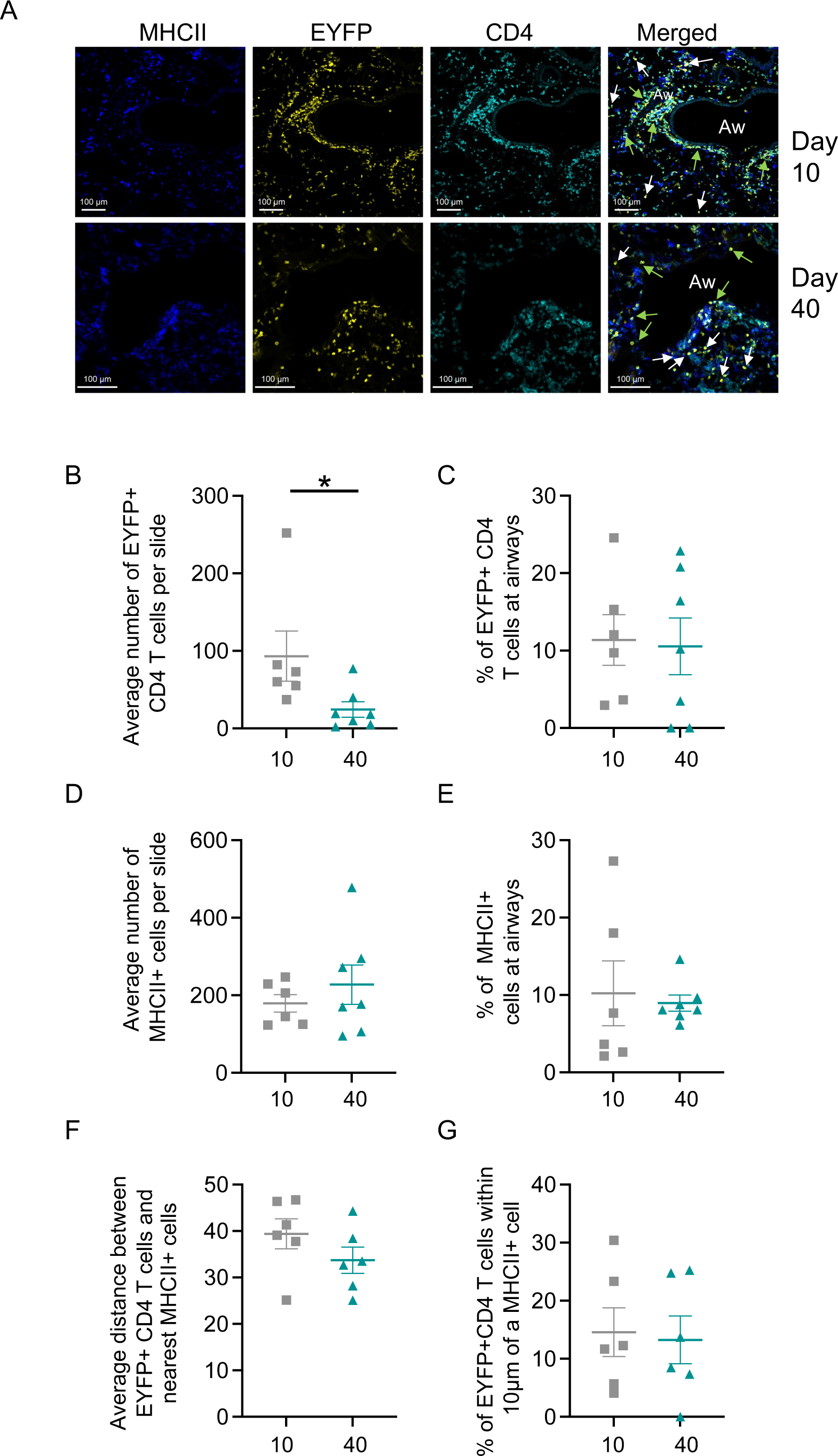
IAV specific CD4 T cells are maintained at lung airways at memory time points. TRACE mice were infected with IAV and lungs taken 10 or 40 days post-infection. Lung sections were stained with the indicted antibodies to identify MHCII+ cells and IAV specific CD4 T cells. In A, images are representative of 6-7 mice per time point from 2 independent experiments and green and white arrows show EYFP+CD4+ cells near or further away from airways respectively; scale bars are 100mm. In B-G, each symbol represents a mouse and the line shows the mean of the group, error bars are SEM. In B, data are not normally distributed (tested by Shapiro-Wilk) and difference tested between time points tested by a Mann-Whitney U test, *: p<0.05.

We measured the distance between the EYFP+CD4 T cells and the nearest MHCII+ cell to ask whether the likelihood of cell interactions altered between the primary and memory time points. The average distance to the nearest MHCII+ cells was the same at both time points, Figure 4F. We also asked what percentage of the EYFP+CD4+ cells were within 10mm of the nearest MHCII+ cell. The distances measured are from cell centroid to centroid, and thus cells within 10mm or less of each other are likely to be touching and potentially interacting. At primary and memory time points, a mean of 11% and 13% respectively of the EYFP+CD4+ cells were within 10mm of an MHCII+ cell, Figure 4G. Together, these data suggest that IAV specific T cells maintain similar levels of contact with APCs at the primary and memory time points.

### 3.4 Immunodominant but not polyclonal IAV-specific memory CD4 T cells have a reduced IFNg response following TCR-MHC blockade

Our imaging studies indicated that lung IAV-specific CD4 T cells are likely to have multiple interactions with APCs during IAV infection. In mice, IAV is cleared within the first ten days of an infection^30, 31^ but antigen can be presented to CD4 T cells for at least 35 days post-infection^38^. We wanted to investigate whether presentation of antigen in the lung was important in the generation of memory CD4 T cell function and location. To address this, we delivered an MHCII blocking antibody^16^ intranasally at day 6 and 12 post-infection. These time points were chosen to allow an initial T cell response to be primed and to block T cell receptor (TCR) pMHC interactions during the peak response in the lung^29, 37^. Importantly, the antibody treatment did not alter the infection-induced weight loss, Supplementary Figure 3.

First, to confirm that the anti-MHCII bound to the different APC populations in the lung, we used fluorescently labelled antibody and examined the lung MHCII+ populations after 2 hours. Anti-MHCII bound to all lung MHCII+ cells including monocytes, macrophages, B cells and DCs, Supplementary Figure 4A-C. In contrast, an isotype control antibody bound to only a small proportion of cells. Some i.n. instilled anti-MHCII labelled cells could be found in the draining lymph node, however, only a small percentage (5.7+/-7.4) of the MHCII+ cells were labelled, Supplementary Figure 4D.

To investigate the consequences of the anti-MHCII treatment, we used flow cytometry to identify IAV-specific CD4 T cells that recognise the immunodominant nucleoprotein (NP) peptide, 311-325, or by gating on EYFP+ CD4 T cells in infected TRACE mice, Figure 5A-B, gating in Supplementary Figure 5. Our prior study demonstrated similar kinetics and differentiation into cytokine producing cells between CD4 T cells identified via these two methods^19^.

**Figure 5.**
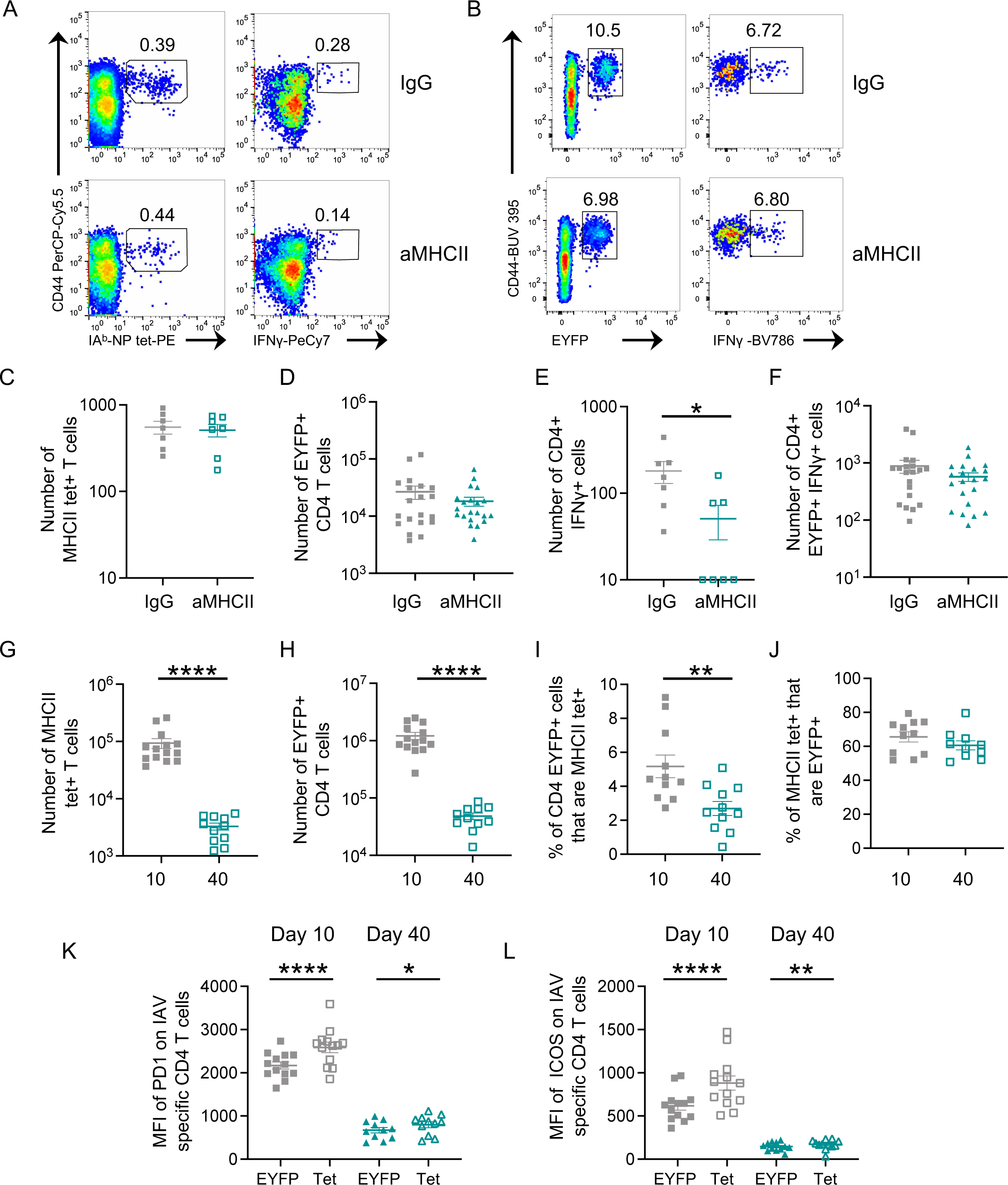
Immunodominant but not polyclonal IAV specific memory CD4 T cell have a reduced IFNg response following anti-MHCII treatment. C57BL/6 (A, C, E) or TRACE (B, D, F) mice were infected with IAV on day 0 and given control IgG or anti-MHCII intranasally on days 6 and 12. On day 40, IAV specific CD4 T cells were detected *ex vivo* using IA^b^/NP_311-325_ tetramers (C) or following restimulation with NP_311-325_ peptide (E) or co-culture with IAV-Antigen treated bmDCs (D,F). In G-L, TRACE mice were infected with IAV and CD4+EYFP+ cells examined following 10 or 40 days. In A-B, cells are gated on CD4+ cells as shown in Supplementary Figure 5 and the numbers are the percent of cells in the indicated gate. Data are from 2-5 independent experiments with a total of 7-8 mice/ time point (A, C, E), 20-21 mice (B, D, F) and 10-12 mice (G-L). Symbols show the mean of the group and error bars are SEM. In E, data are not normally distributed (tested by Shapiro-Wilk) and difference tested by Mann-Whitney U test, the Y axis is set at the level of detection. In G, data are not normally distributed and tested via a Mann Whitney U test. In H-L, data are normally distributed and differences tested using a t-test in H-J and paired t-tests between cell types in K-L. In all graphs, *: p<0.05; **:p<0.01; ***: p<0.001; ****p<0.0001.

There were no differences in the numbers of either IA^b^/NP_311-325_ tetramer (tet)+ or EYFP+ CD4 T cells between animals treated with anti-MHCII or IgG control antibody, Figure 5C-D. To examine production of the key T cell derived anti-viral cytokine, interferon (IFN)-g, we restimulated the lung cells *ex vivo* with NP_311-325_ or bone marrow derived dendritic cells (bmDCs) incubated with sonicated IAV antigens^19, 21^. While anti-MHCII treatment reduced the number of NP_311-325_ specific IFNg+ CD4 T cells within the memory pool, there was no difference in the number of EYFP+ IFNg+ in infected TRACE mice, Figure 5E-F. These data suggest there may be different requirements for sustained antigen presentation in the formation of memory CD4 T cells specific for different IAV epitopes.

### 3.5 Lung immunodominant IAV-specific CD4 T cells express higher levels of T cell activation markers than polyclonal IAV-specific T cells

To explore the difference in dependency on TCR-pMHCII interactions on the generation of IFNg+ memory cells, we examined the phenotype of these cells in IAV infected TRACE mice. As expected, the numbers of both CD4 T cells detected by MHCII tetramer and EYFP+ CD4 T cells declined from day 10 to day 40 post IAV infection, Figure 5G-H. When we examined the percentages of EYFP+ cells that were also MHC-tet+, we found this declined from day 10 to day 30 post-infection, Figure 5I. These data indicate that CD4 T cells specific for NP_311-325_ may be less likely to enter the memory pool than other IAV specific CD4 T cells. In contrast, the percentages of MHC-tet+ cells that were also EYFP+ were consistent at both time points, Figure 5J.

We examined expression of PD1 and ICOS, two molecules associated with T cell activation, and that are expressed by some memory T cells^13, 14, 19, 39^. As expected, memory CD4 T cells expressed lower levels of both molecules than cells at day 10 post-infection. However, at both time points, MHCII tet+ CD4 T cells expressed higher levels of PD1 and ICOS in comparison to the remainder of the EYFP+ population, Figure 5K-L and Supplementary Figure 6. We found a similar pattern of expression in CD4 T cells from the lung draining lymph node and spleen at day 10 post-infection but the difference at the memory time point was minimal or absent, Supplementary Figure 7. Together these data suggest that, particularly in the lung, CD4 T cells specific for an immunodominant IAV epitope express higher levels of molecules associated with T cell activation than T cells of other IAV-specific CD4 T cells. This distinction may explain their reduced entry into the memory pool and greater dependence on antigen presentation compared to other IAV-specific CD4 T cells.

### 3.6 Anti-MHCII treatment alters the proportion of memory IAV-specific CD4 T cells at the airways

To investigate whether the location of IAV-specific memory CD4 T cells was altered following anti-MHCII treatment, we examined lung sections by immunofluorescence at day 40 post-infection. As in the flow cytometry experiments, mice were treated with anti-MHCII or control IgG intranasally at day 6 and 12 post-infection. It is not possible to use MHCII tetramers on frozen sections. However, we examined the location of MHCII+ cells and EYFP+ CD4 T cells and determined the expression of PD1 by the CD4 T cells, Figure 6A.

**Figure 6:**
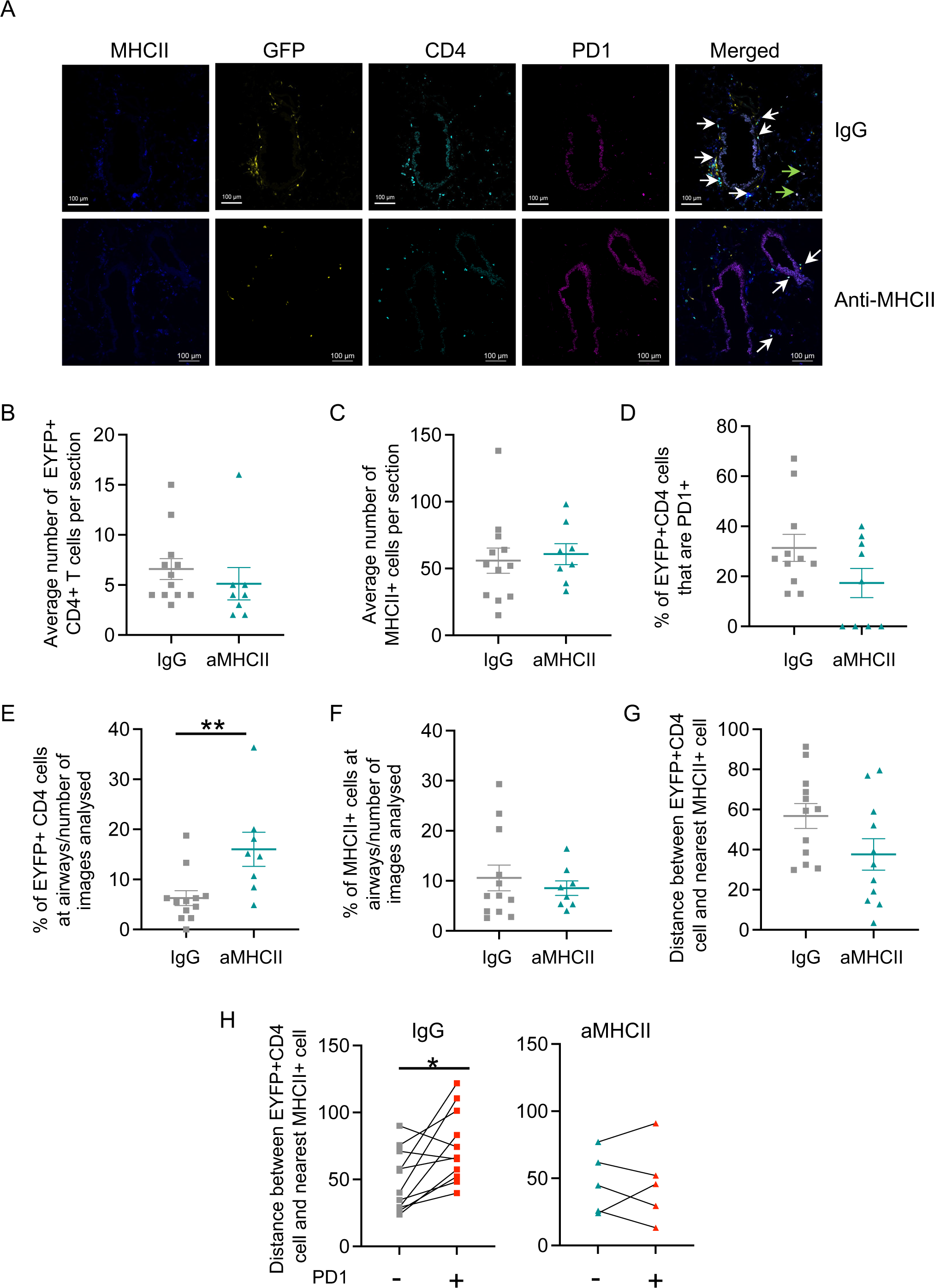
Anti-MHCII treatment alters the location of IAV-specific CD4 T cells. TRACE mice were infected with IAV on day 0 and on day 6 and 12 given control IgG or anti-MHCII intranasally. On day 40, lungs were frozen and tissue sections stained with the indicated antibodies. Data are combined from two independent experiments with 7-12 mice per group. In A, white and green arrows indicate EYFP+CD4+ cells and EYFP+CD4+PD1+ cells respectively; the scale bar is 100mm. In B-G, each symbol represents a mouse, the line shows the mean of the group and error bars are SEM. For each mouse 2-5 sections were analysed. In B, D, E and F data are not normally distributed (tested by Shapiro-Wilk) and differences tested by Mann Whitney U test. In C, G and H, data are normally distributed and differences tested by t-test (C,G) or Paired t-test (H). In H, 4 anti-MHCII mice are removed as no PD1+ cells were found in the analysed sections.

In line with our flow cytometry data, neither the numbers of EYFP+CD4 T cells nor MHCII+ cells identified per slide were altered by the antibody treatment, Figure 6B-C. There was a slight but non-significant reduction in the percent of EYFP+ CD4 T cells that were PD1 positive and in some anti-MHCII treated mice, we could not detect any PD1+ EYFP+ CD4 T cells, Figure 6D.

To investigate whether the anti-MHCII treatment affected CD4 T cell location, we assessed whether the EYFP+ CD4 T cells were proximal to airways. We normalised for the number of slides with airways by dividing the percentages of EYFP+CD4 cells at airways by the numbers of slides that had airways. These data show that blocking MHCII-TCR interactions during the primary immune response led to an increase of memory CD4 T cells at airways, Figure 6E. There was not, however, any impact on the location of MHCII+ cells themselves, Figure 6F.

We also measured the distances between the EYFP+ CD4 T cells and the nearest MHCII+ cell. Anti-MHCII treatment led to a slight but non-significant decline in the distance between the EYFP+ CD4 T cells and nearest MHCII+ cell, Figure 6G. More strikingly, there was a clear difference in the distance between EYFP+CD4 T cells and the nearest MHCII+ cell depending on the T cell’s expression of PD1. EYFP+ CD4 T cells that were PD1+ were further away from MHCII+ cells than PD1 negative T cells, Figure 6H. These data suggest that the PD1 expression might not be a consequence of a recent interaction with an APC. In contrast, in anti-MHCII treated animals in which we could detect PD1+ cells, this distinction in distance to the nearest MHCII+ cell was lost.

Taken together with the flow cytometry results, these data suggest that interactions between CD4 T cells in the lung during the later stages of the primary response are required to shape important attributes of memory CD4 T cells. Antigen-driven interactions may be required for optimal cytokine responses by some T cells. In contrast, these same interactions may reduce the opportunity for the CD4 T cells to position themselves near airways, often the initial infection site for respiratory viruses.

## 4. Discussion

Heterogeneity within the memory CD4 T cell pool has focussed on cell phenotype, gene expression, and location^40, 41^. Memory cell location has mainly been examined at the organ level, comparing cells within lymphoid and non-lymphoid tissues. Here we focused on the micro-location of lung IAV-specific memory CD4 T cells determining how the location of these cells alters from the primary to the memory phase of the response. Our data demonstrate that both primary and memory CD4 T cells are found in multiple micro-locations including at airways and in packed clusters with other immune cells. Importantly, our data show that local TCR-pMHC signals delivered during the later stages of the primary immune response are not required for the formation of memory cells in the lung. However, these signals do influence the ability of immunodominant CD4 T cells to produce IFNg and limit the access of memory CD4 T cells to airways.

We analysed IAV-driven changes to the lung using H&E and confocal analysis of immune cells. In both cases, our data demonstrate persistent changes to the landscape of the lung following infection. Clusters of immune cells that persist for months following IAV infection have been documented previously^22–25, 28^. The clusters in our studies do not have clear B and T cell zones and thus we have not referred to them as induced bronchus-associated lymphoid (iBALT) tissue or tertiary lymphoid organs^26, 27^. We found a wide range of immune cells within clusters including those expressing B220, CD64 and CD11c. The B220+ cells are likely B cells, known to be retained in the lung and important for protection to re-infection, in particular following challenge with the same IAV strain^24^. The IAV specific CD4 T cells contained within clusters with the B220+ cells are likely to overlap with tissue resident helper cells that can promote protection from a different IAV strain^13, 14^.

IAV-specific CD4 T cells were found in close proximity to MHCII+ cells at both primary and memory time points following infection. These data suggest that the CD4 T cells may continue to interact with APC despite viral clearance. IAV antigen can be maintained in the lung and/or lung draining lymph nodes for weeks-months following infection^38, 42^ and these interactions may shape the survival and subsequent function of memory CD4 T cells.

We found that IAV-specific CD4 T cells were maintained at airways from the primary to the memory response. Moreover, we increased the proportion of IAV-specific CD4 T cells at airways by treating the mice with a blocking anti-MHCII antibody during the late primary response. These data indicate that interactions between CD4 T cells and APCs at this time shape the subsequent memory pool. Lung airway epithelial cells are some of the first cells to be infected with IAV^43^ and by positioning themselves near airways, memory CD4 T cells have the potential to respond rapidly following a re-infection, either in an antigen-specific manner via presentation by an APC, including epithelial cells^44, 45^, or potentially, in response to local cytokines^46^.

It is unclear why interactions with lung APCs would limit airway memory CD4 T cells. Potentially sustained interactions drive increased effector cell differentiation and that these highly differentiated cells present a risk of immune-driven damage at delicate airways^47, 48^. This hypothesis would fit with our finding that anti-MHCII treatment reduced IFNg production by immunodominant, but not all, IAV-specific memory CD4 T cells. It is likely that the immunodominant T cells have proliferated substantially during the infection, having received strong activation signals, corresponding with our finding that they express higher levels of PD1 and ICOS than the majority of IAV-specific CD4 T cells. Interestingly, lung IAV specific memory CD4 T cells expressing PD1 were more likely to be further away from an MHCII+ cells than those with low or no PD1 expression. These data suggest that cells which receive strong activation signals may be restricted from future interactions by limiting their access to MHCII APCs. An alternative explanation is that PD1 expression indicates cells that have recently received a TCR signal and that these cells have migrated away from the APC following a productive interaction.

Together, these data suggest that interactions between lung CD4 T cells and APCs temper, but are not required for, the generation of memory cells. These data highlight the value of analysing responding T cells using multiple readouts to provide a more complete understanding of the molecular regulation of memory cell formation. As the anti-MHCII antibody bound to many types of cells and the IAV-specific CD4 T cells could be found near to multiple MHCII+ cells, it was not possible to identify which APCs may be involved in tempering the generation of memory CD4 T cells. Potentially, all the different MHCII+ cells could influence memory CD4 T cell generation.

In summary, this study extends our understanding of heterogeneity within the lung memory CD4 T cell pool. CD4 T cells within and outside lung immune clusters will experience different signals and interact with different cell types suggesting distinct memory niches that may influence the type and function of lung memory CD4 T cells^49^. It will be key for future studies to understand the dynamics of cells within these different environments. For example, to address whether cells can move into and out of clusters, what controls these decisions during the maintenance of immune memory during, and, importantly, how these cells respond to successive infections.

## Data availability

All raw data are available upon request to the corresponding author.

## Competing interests

The authors have no competing interest.

## Acknowledgments

We thank the staff within the School of Infection and Immunity Flow Cytometry Facility, the Glasgow Imaging Facility, and Biological Services at the University of Glasgow for technical assistance. We thank Pablo Murcia for helpful conversations on viral responses. We thank the NIH tetramer core facility for the provision of IA^b^-NP_311-325_ tetramers. The work was supported by a Marie Curie Fellowship (334430), a Wellcome Trust Investigator Award (210703/Z/18/Z) to MKLM, and a Rosetrees’ Trust Seedcorn Grant (Seedcorn2020\100017) awarded to JCW.

## Author contributions

KEH: conception, investigation, formal analysis, visualization, data curation, writing -original draft and reviewing/editing. JCW: funding acquisition, investigation, formal analysis, visualization, data curation, writing original draft – reviewing and editing. CP: investigation, formal analysis, writing – reviewing and editing. EB: investigation, formal analysis, reviewing and editing. AM: investigation, formal analysis, reviewing and editing. JIG: investigation, formal analysis, writing - reviewing and editing. TP: investigation, writing – reviewing and editing. EWR: formal analysis, writing -reviewing and editing. MKLM: Project management, funding acquisition, conception, investigation, formal analysis, visualization, writing -original draft and reviewing.

## Figure Legends

**Supplementary Figure 1.**
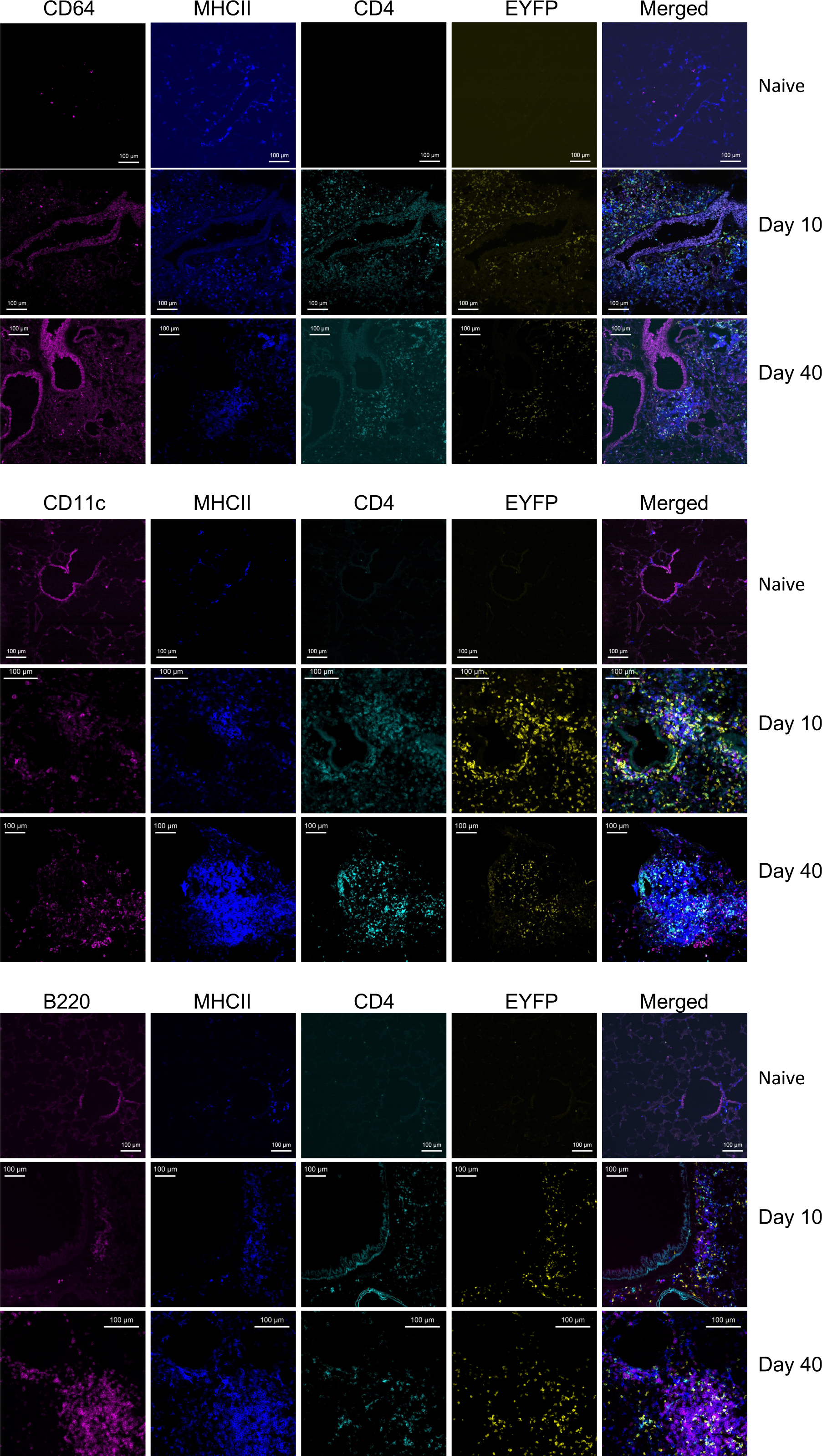
Identification of MHCII+ cells in IAV infected lungs at primary and memory timepoints. TRACE mice were either naïve or infected with IAV and lungs removed 10 or 40 days post-infection. Lungs were sectioned and stained with the indicated antibodies to identify IAV specific CD4 T cells and MHCII+ CD64, CD11c or B220+ cells. Data are representative 3-5 mice per time point and scale bars are 100μm.

**Supplementary Figure 2.**
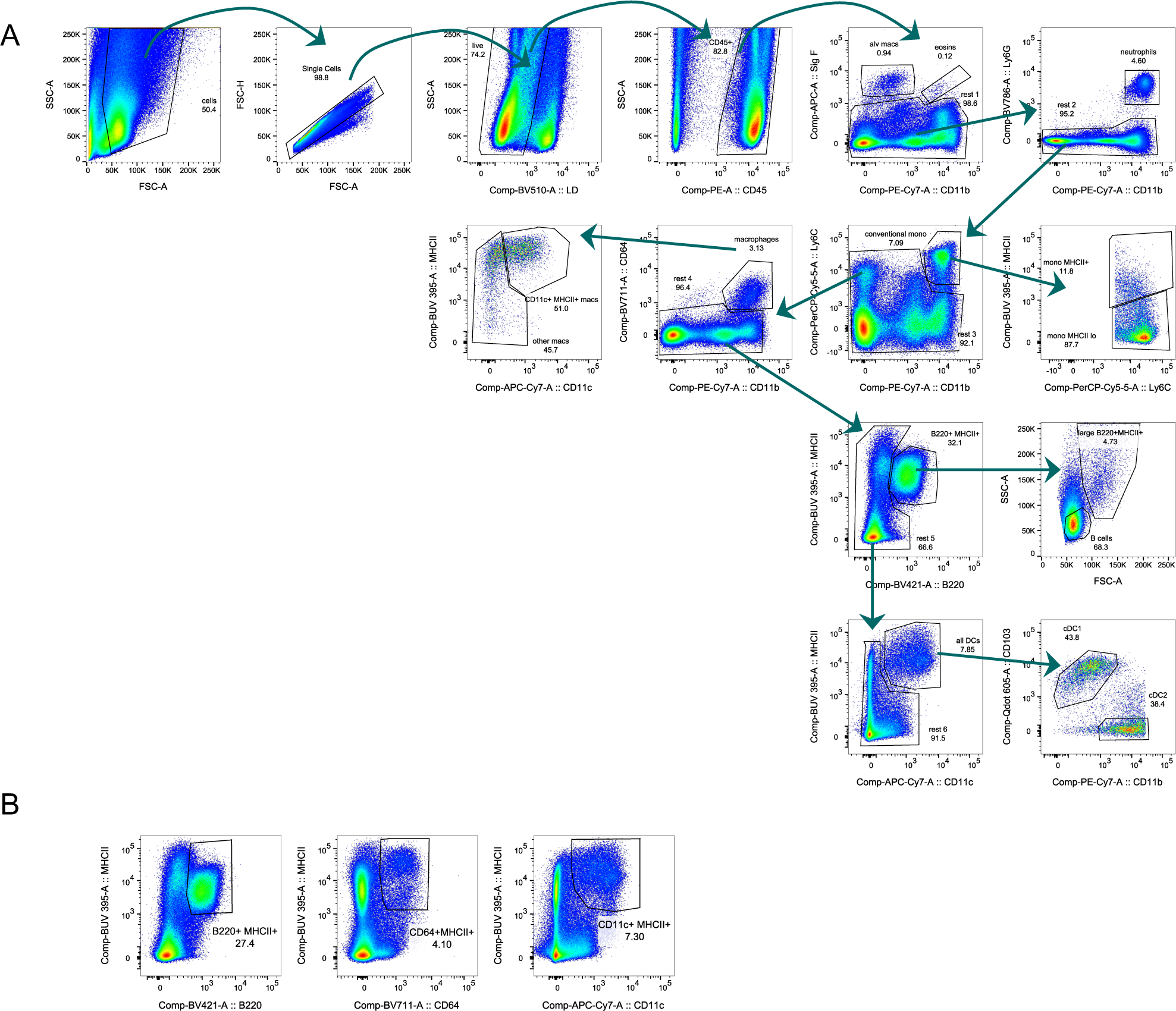
Gating strategy to identify MHCII+ populations. Flow cytometry analysis of lungs from a mouse infected with IAV 30 days earlier. A shows example gating. In B, cells from the same sample are first gated through the CD45+ gate in A to identify populations of MHCII+ B220, CD64 and CD11c+ cells.

**Supplementary Figure 3.**
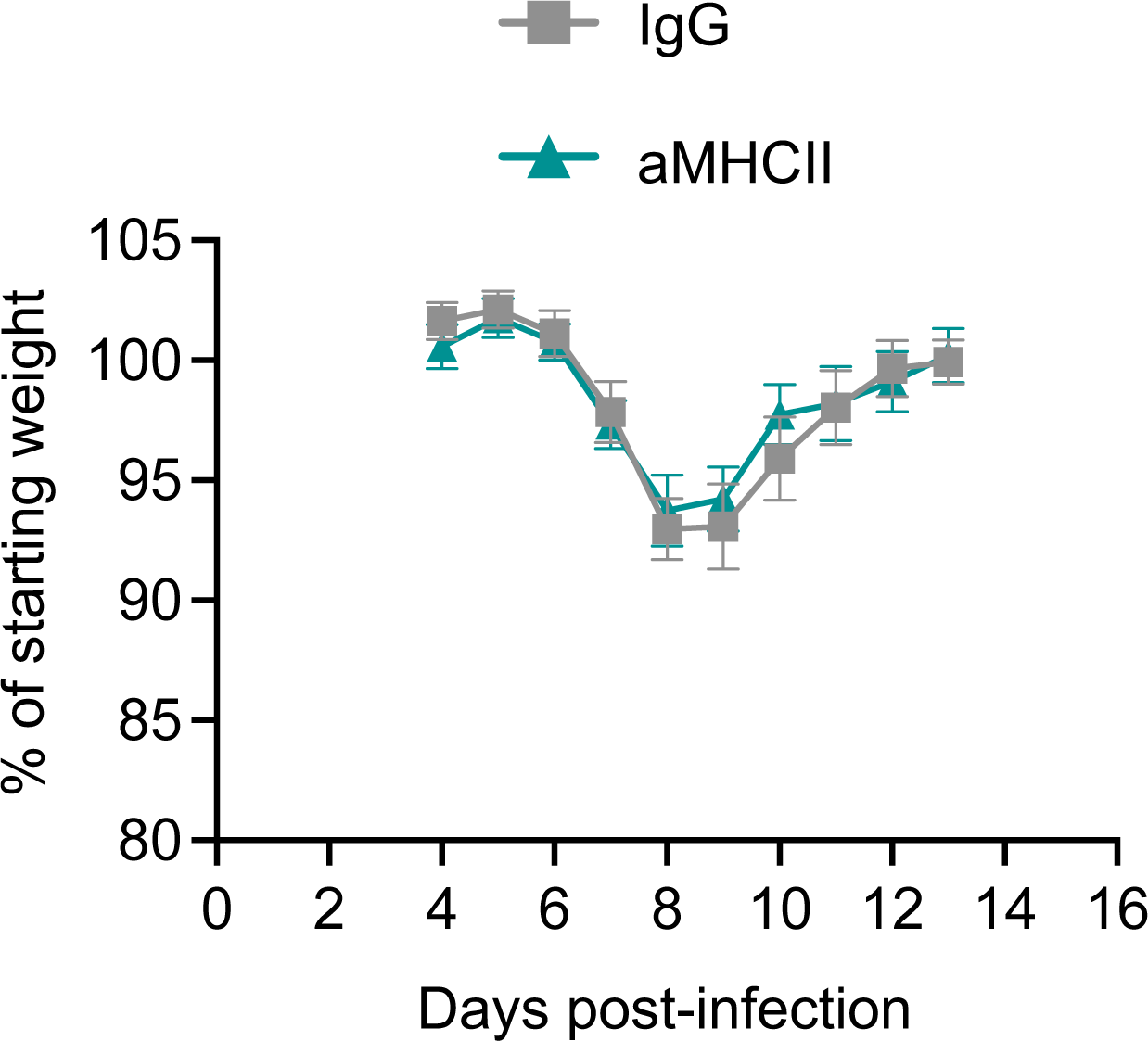
Anti-MHCII treatment does not alter IAV-induced weight loss. TRACE mice were infected with IAV on day 0 and weighed from day 4 until day 14 post-infection. Mice received 100μg of either control IgG or anti-MHCII intranasally on day 6 and 12 post-infection. Data are combined from 6 experiments with a total of 21-22 animals per group.

**Supplementary Figure 4.**
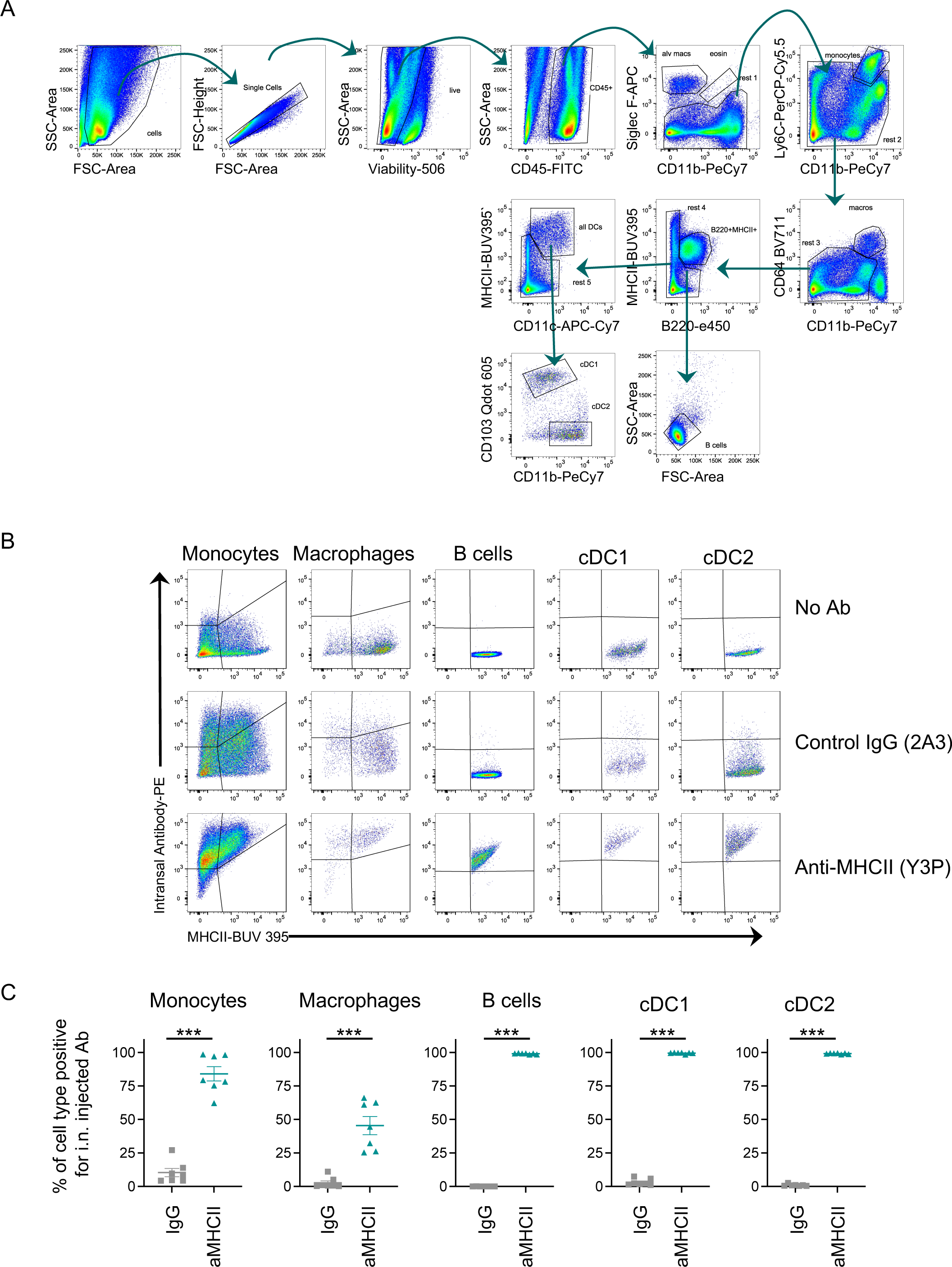
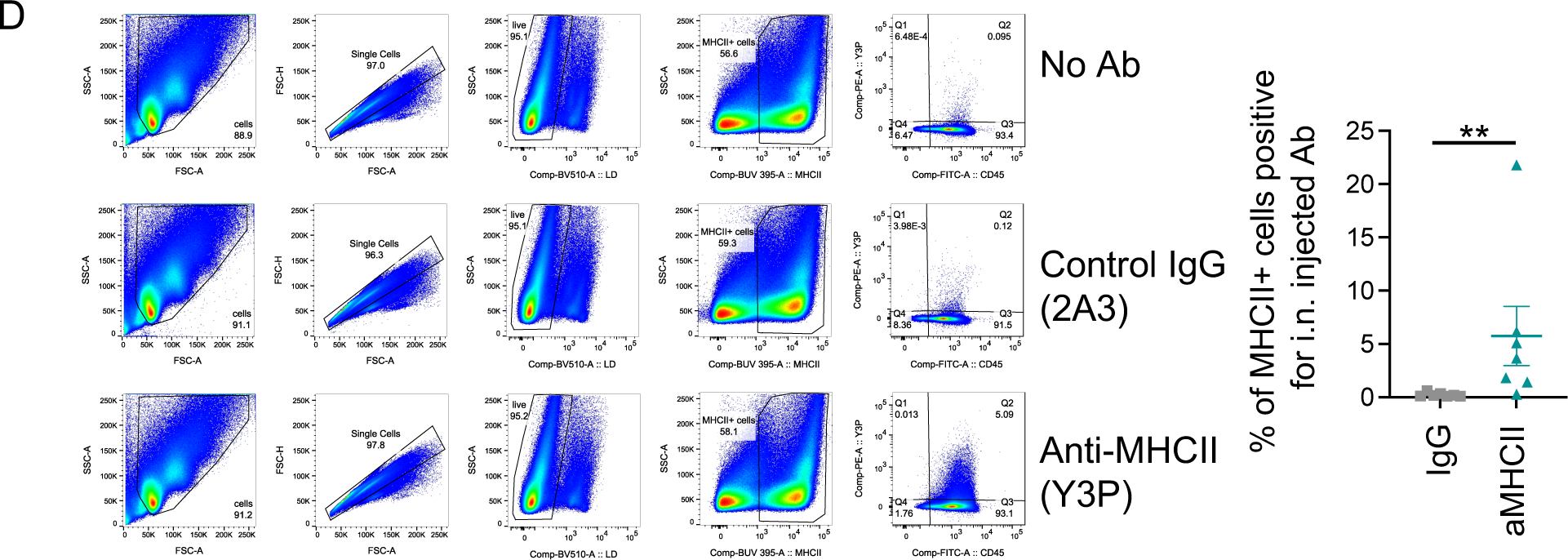
Intranasal anti-MHCII binds to MHCII+ cells in the lung but to few MHCII+ cells in the draining lymph node. C57BL/6 mice were infected with IAV i.n. on day 0 and received 100μg of either control IgG or anti-MHCII labelled with Alexa-Fluor546 i.n on day 6. 2 hours later, single cell suspensions of lung (A-C) and mediastinal lymph node cells (D) were stained for flow cytometry. The gating strategy in A was used to identify lung monocytes, macrophages, B cells, cDC1 and cDC2. The data are representative of two experiments with a total of 8 (control IgG) and 7 (anti-MHCII) mice. In C and D, the percentages of the cell populations double positive for the i.n. antibody and anti-MHCII labelled *ex vivo* are shown, error bars are SEM and the horizontal line shows the mean of the group. Data were not normally distributed (tested by Shapiro-Wilk), significance tested with a Mann Whitney U test, **:p<0.01 ***: p<0.001.

**Supplementary Figure 5.**
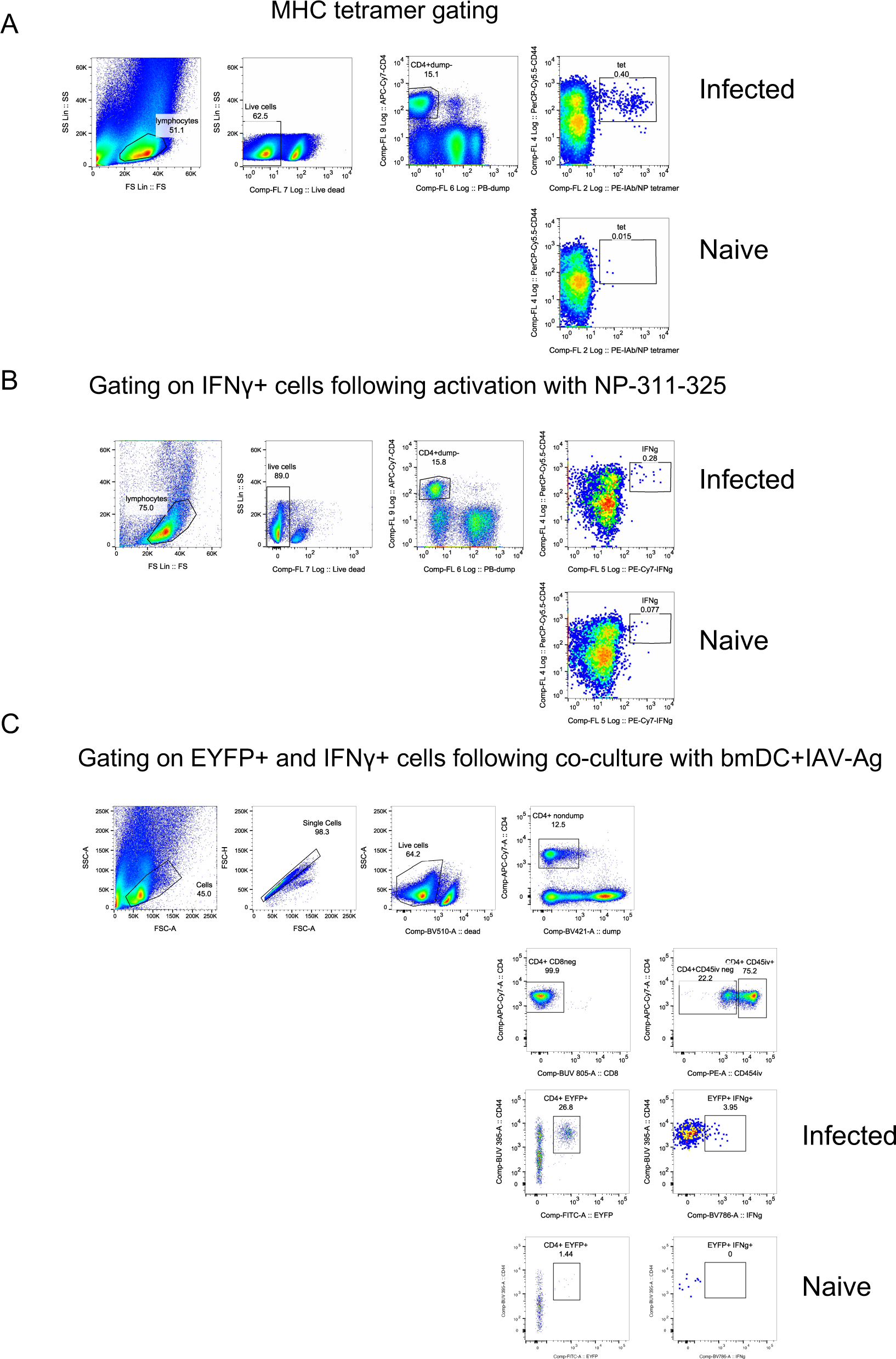
Gating to identify MHC tetramer+ and EYFP+ T cells IAV specific CD4 T cells C57BL/6 mice (A, B) or reporter. TRACE mice (C) were infected with IAV i.n. on day 0 and received 100μg of either control IgG or anti-MHCII i.n on day 6 and 12. On day 40, single cell suspensions of lung cells were stained for flow cytometry either directly *ex vivo* (A), following 6 hours of *ex vivo* restimulation with either NP_311-325_ peptide (B), or bone marrow derived DCs (bmDCs) incubated with IAV-Ag (C). All *ex vivo* restimulations were done in the presence of Golgi Plug. In A-B, mice were perfused with PBS-EDTA to remove cells within the blood; in C, blood cells were labelled with PE-labelled CD45 injected i.v. 3 minutes prior to euthanasia.

**Supplementary Figure 6.**
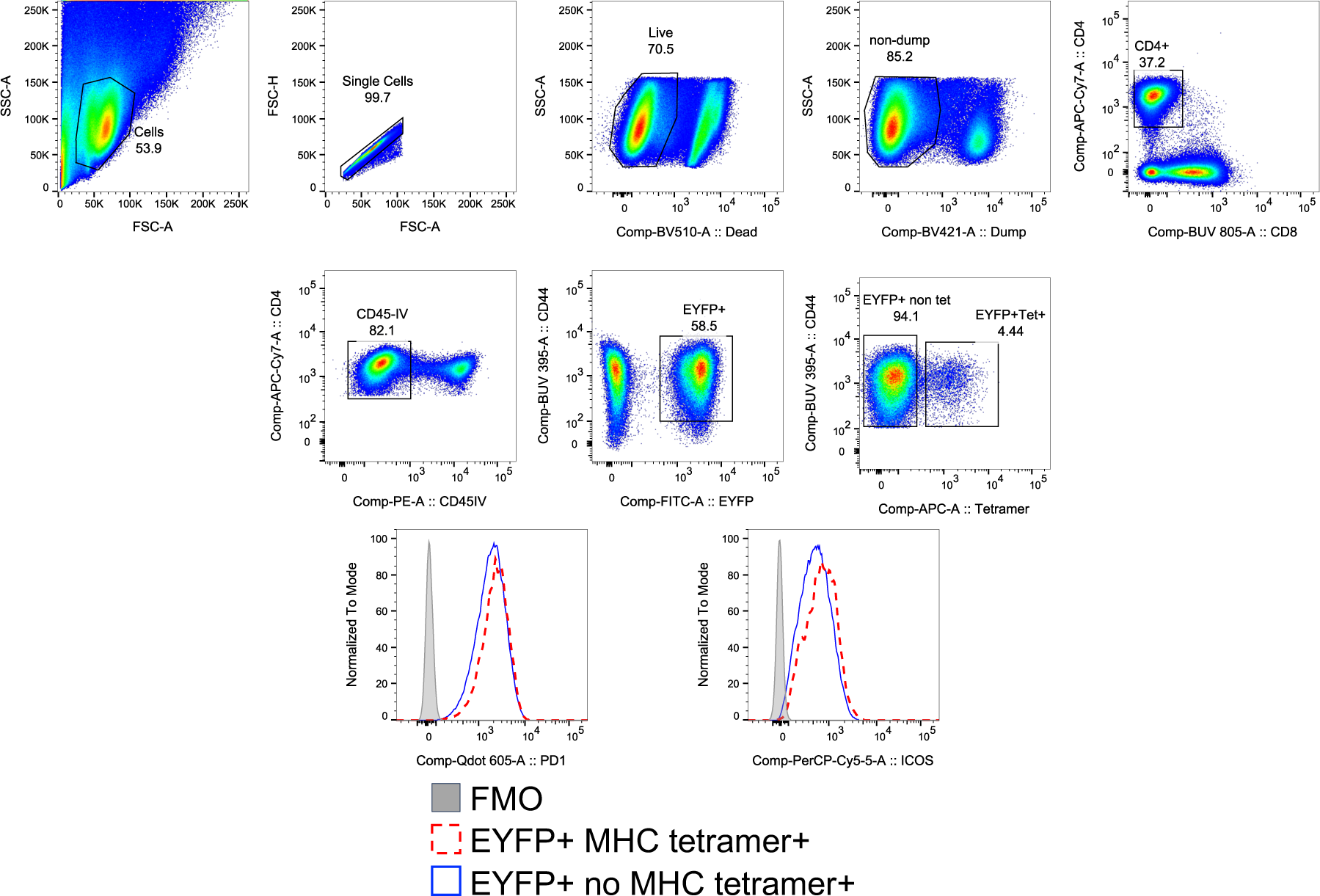
Example gating for PD1 and ICOS on IAV specific CD4 T cells identified by EYFP+ expression on TRACE mice or MHCII tetramers. TRACE mice were infected i.n. with IAV on day 0 and 10 days later injected with CD45 PE i.v. 3minutes prior to removal of tissues. Single cell suspensions of the lung were stained with MHCII tetramer and surface antibodies and cells examined by flow cytometry. Cells are gated as shown and the numbers indicate the percentages of cells within the gates. Data are from the experiment shown in Figure 5.

**Supplementary Figure 7.**
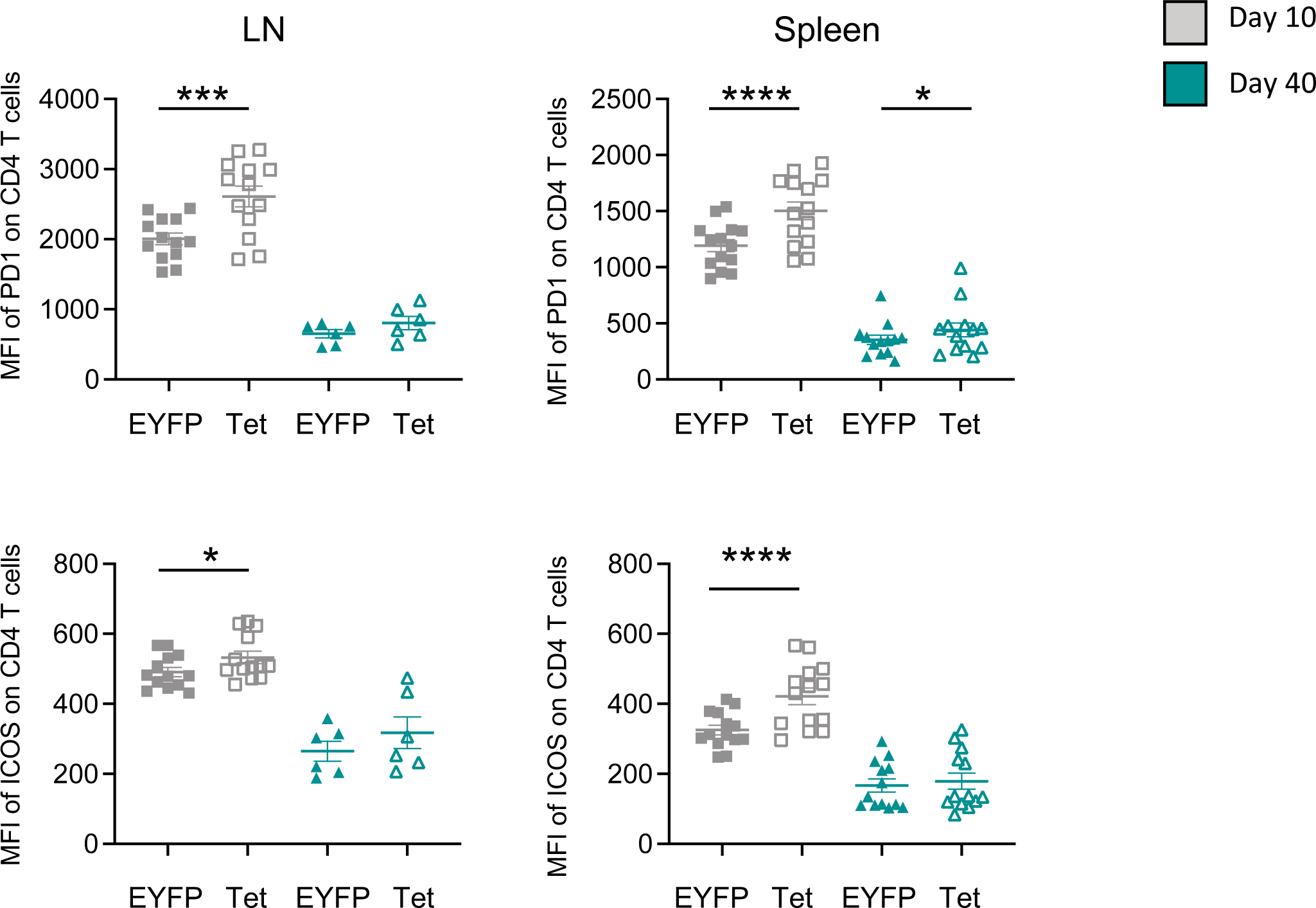
MHC tetramer+ CD4 IAV specific T cells in the spleen and lymph node express higher levels of PD1 and ICOS than EYFP+ cells at day 10 but not day 40 post-infection. TRACE mice were infected i.n. with IAV on day 0, 10 or 40 days later these mice were injected with CD45 PE i.v. 3minutes prior to removal of tissues. Single cell suspensions of the spleen and draining lymph node were stained with MHCII tetramer, surface antibodies and viability dye and cells examined by flow cytometry. Cells are gated as shown in Supplementary Figure 5. Data are from two experiments with 5-8 mice per group. 2 lymph node samples were excluded from the first memory experiment and 5 from the second as less than 10 MHCII tetramer+ cells were present in the FACS plots. Error bars are SEM and differences between EYFP+ and MHCII tet+ cells tested using paired T-tests. *:p<0.05; ***:p<0.001; ****:p<0.0001.

## Notes

### Competing Interest Statement

The authors have declared no competing interest.

### Summary of Updates

Small changes to figure 1 to add additional H&E staining and some small corrections of the text.

